# Simple and low-cost amplifier system for the study of a ventral nerve action potential in earthworm

**DOI:** 10.1101/832055

**Authors:** Orlando Jorquera

**Affiliations:** Formation Center in Environmental Science (CFCAm), Campus Sosigenes Costa, Porto Seguro, Federal University of Southern Bahia-UFSB, Rodovia Porto Seguro-Eunapolis, BR-367, km 10, CEP 45810-000, Brazil

**Keywords:** Action potential, giant ventral nerve cord, earthworm, low-cost amplifier circuit

## Abstract

Basic Biophysics or Neurobiology courses encounter problems when starting their practices due to the complexity and high cost of the equipment. As a consequence, these experiences are replaced by simulation software, leading students to boredom and disinterest, and finally to the lack of understanding of the basic principles of the phenomenon observed. Classical nerve conduction studies for the visualization of action potentials are an example of this, being able to be developed with simple and low-cost amplification circuits, as described in this article. The system showed stability, expected amplification and high signal-to-noise ratio. The action potential of the medial giant fiber (MGF) as lateral giant fiber (LGF) in the ventral nerve cord of the earthworms *Lumbricus terrestris* was recorded. The experimental system has met expectations and regains practical, effective and stimulating learning for students.

## 1. Introduction

In the study in classical neurophysiology of neural conduction mechanisms, must pass by the reading of the works carried out by Hodgkin-Huxley. They describe an electrical change recorded in nerve fibers known as action potentials (AP) [1]. Because most of the nerve fibers are extremely small, AP records were initially performed on giant squid axons (*Loligo forbesi*), but obtaining them is difficult because of their geographic and seasonal availability, leading to the search for other animals with the same feature.

In general, invertebrates, such as earthworms, have larger nerve cords than other taxonomic groups, facilitating conduction studies in the nervous system,[2],[3]. These one have a ventral nerve cord large enough to be used in classical electrophysiology studies [2],[4],[5],[6],[7],[8],[9].

On the other hand, most AP records published are made with high-cost amplification equipment, making it difficult to access these types of practical studies by replacing them with virtual experiments through software, causing students to feel bored and disinterested in difficult to learn. Nowadays it is possible to find electronic components of low cost for the implementation of this type of amplifier. Traditional action potential recording systems require basic electronic equipment such as an oscilloscope, a stimulator (eg grass stimulator), an amplifier, a low-pass filter, and a data acquisition system. Also, we need a suction electrode and a micro-manipulator. All of this high-cost equipment except the suction electrode.

Recent efforts by some authors [8],[7], in the earthworm AP record, used relatively inexpensive equipment with invasive or non-invasive systems and with or without electrical stimulation, obtaining records without good resolution and with associated noise. The complete setup configuration is important for good records and it is difficult to dispense equipment such as oscilloscope or acquisition systems with an adequate sampling rate.

In an attempt to solve some of these problems the objective of this work was to describe a simple and low-cost amplification circuit for the reproduction of classical studies of action potentials in the ventral nerve cord in earthworms.

## 2. Materials and Methods

### 2.1 Biological material, ventral nerve dissection and experimental system

The procedure was an adaptation as described by Roberts (1962). Earthworms of the species *Lombricus terrestris* were obtained around the Sosígenes Costa Campus of the Federal University of Southern Bahia, Porto Seguro, Bahia, Brazil. For maintenance, they were placed in a container containing soil and minced banana. To dissect the ventral nerve cord the earthworm was anesthetized in 10% alcohol for 6 minutes, washed with water to remove alcohol residue and placed in a petri dish filled with hardened silicone. The worm was placed on the petri dish with the dorsal side up and pinned to 2/3 of the cephalic part. Using a low-cost common magnifying glass to amplify the tissue, the dorsal part was cut with scissors being careful not to damage the intestine and nerves and it was fixed with pins, exposing the digestive system which was removed with tweezers to expose the ventral nerve cord. Saline solution for earthworm [11] consisting of NaCl (6 g/L), KCl (0.12 g/L) CaCl_2_ (0.2 g/L) and NaHCO_3_ (0.1 g/L) was added to wipe scraps of tissue and keep saline medium. The nerve was cut at the bottom with scissors and separated from the tissues by the tip of a suction electrode.

The loose nerve was suctioned with a glass suction electrode consisting of an internal Ag/AgCl electrode and an Ag/AgCl outer coiled to the 1 ml pipette tip, attached to a 30 cm tube and a 5 mL syringe similar to described by Johnson et al., (2007). To chloridizing the silver wire it was placed in a 5% solution of sodium hypochlorite overnight. This suction electrode is attached to a low-cost manipulation system consisting of articulated metal bars and not to a micro-manipulator as commonly used and connected to the amplifier as described in the following section. The stimulus electrodes were connected in the cephalic part of the earthworm with two electrodes the anode (+) closest to the head and the cathode (-) following it (see Fig.1). The preparation was earthed between the first pair of stimulus electrodes and the nerve cord. The action potentials were amplified and visualized using a Tektronix TBS 1072B digital oscilloscope. The computer was running on battery power to reduce electrical noise. The experiments were carried out at an ambient temperature of 28^*◦*^C. To determine the conduction velocity (CV) the following equation was used:

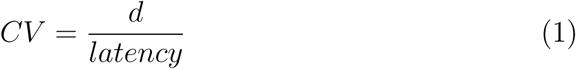

**Fig.1.**
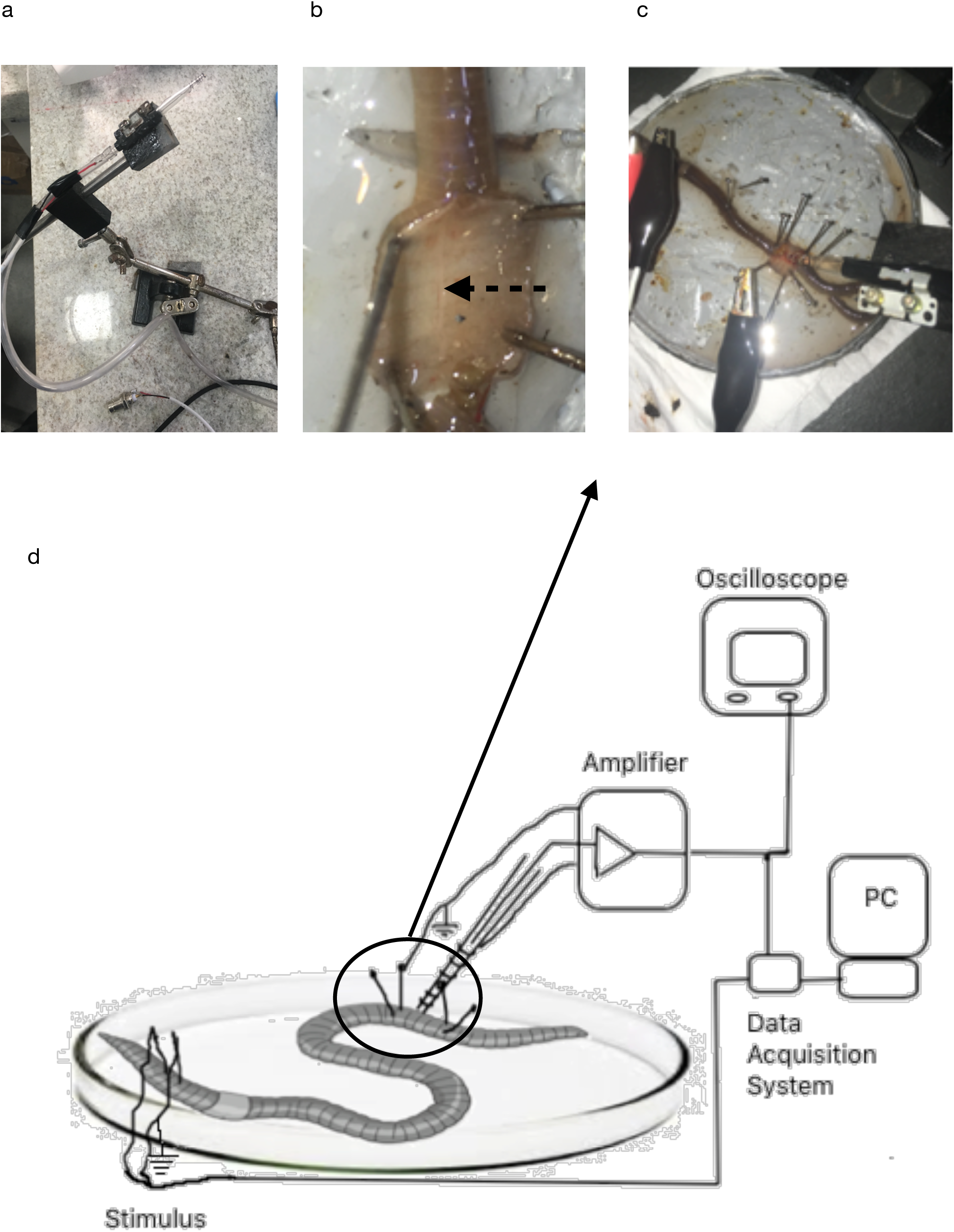
Experimental set up for recording action potentials from a ventral neural cord in the earthworm. a) Suction electrode coupled to the manipulator; b) ventral nerve after dissection (segmented arrow); c) positioning of the stimulation electrodes on the earthworm head, the recording electrode (suction pipette) and the reference electrode near the latter; d) experimental system diagram indicating the animal used the stimulation, amplification, visualization (oscilloscope) and acquisition and recording system (USB-6218, computer).

Where d is the distance between the stimulation electrodes and the suction electrode (5 cm approx.) and latency, which is the time between the stimulus and the first deflection from baseline.

### 2.2 Electronic circuit, stimulus system and data acquisition

To amplify the voltage of action potential was made a circuit consisting of two followers amplifier connected at the input of a differential amplifier (INA128P) and an offset regulator made with an amplifier type TL071CN and two 9 volt batteries that also serve to power the circuit (Fig.2). This offset regulator is for adjusting the baseline close to 0 mV. The circuit simulation can be seen in the following URL, http://tinyurl.com/yy8vuz49. According to the configuration of the suction electrode connections, the internal electrode was connected to the non-inverting input and the external electrode (screwed into the suction electrode) was connected to the inverting input of the differential amplifier INA128. The output of the differential amplifier was connected to an inverted offset regulator amplifier TL071CN which gives finally a non-inverted signal for inverting input and inverted signal for non-inverting input. In the case of the experimental system assembled, first detects the potential variation in the non-inverting input (internal suction electrode) compared to the inverting input (external suction electrode) which gives the expected response to an extracellular register.

**Fig.2.**
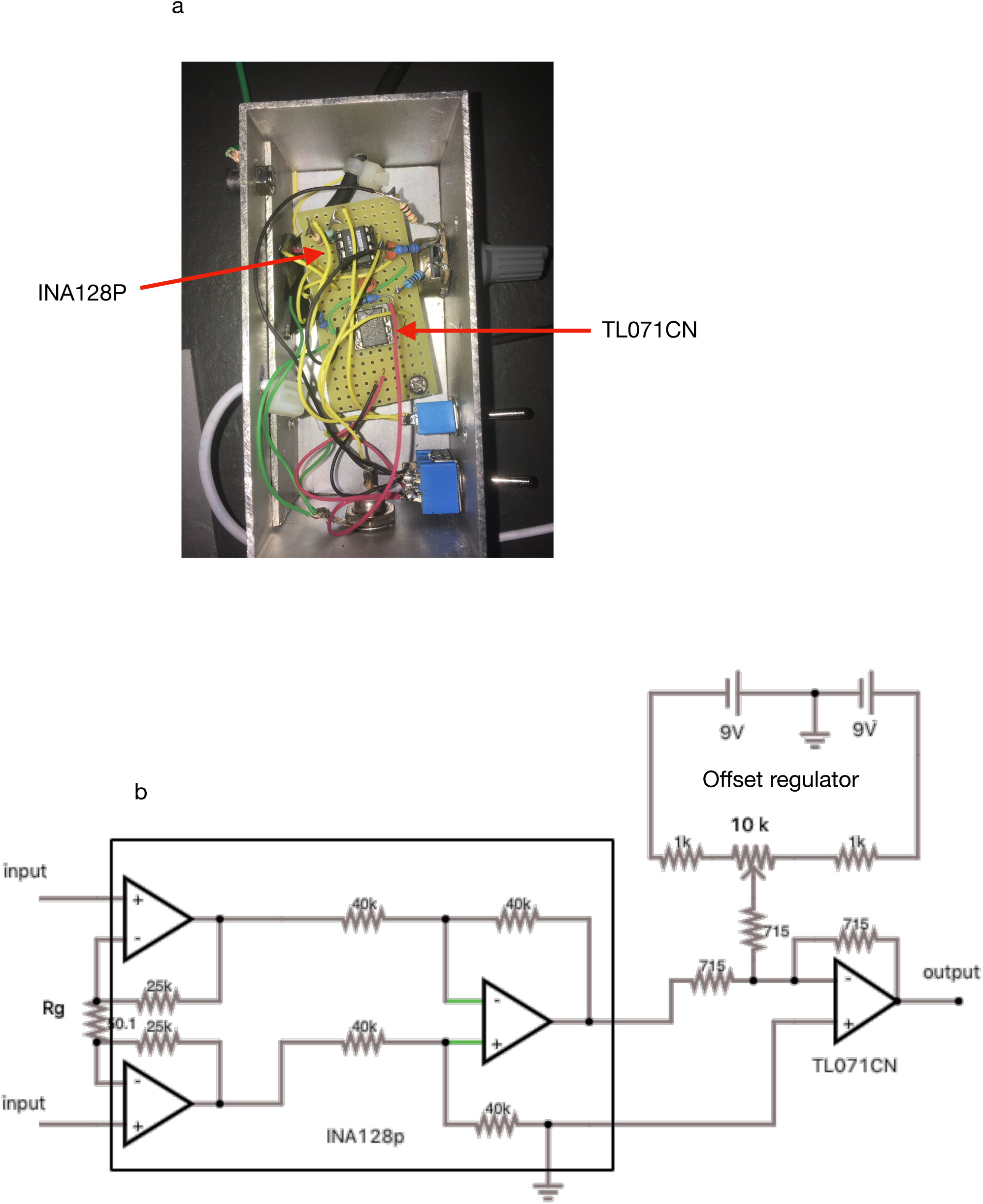
Low-cost amplifier for recording action potentials. a) Photograph of the assembled system; b) circuit diagram used for mounting the amplifier consisting of two operational amplifiers, one INA128p, and one TL071CN plus one offset regulator.

Table 1 shows the components used to mount the amplifier and the approximate cost for each of them.

**Table 1:** Material needed to build the amplifier circuit and associated cost. Value quoted in Digi-key electronics (www.digikey.com).

| Component | Quantity | Cost US\$ | Total (US\$) |
| --- | --- | --- | --- |
| INA128p | 1 | 11.16 | 11.06 |
| TL071 | 1 | 0.4 | 0.4 |
| Resistor 511 ohm | 1 | 0.11 | 0.11 |
| Resistor 47 ohm | 1 | 0.1 | 0.1 |
| Resistor 1kohm | 1 | 0.12 | 0.12 |
| Resistor 715 ohm | 3 | 0.18 | 0.54 |
| Capacitor 0.1 uF | 2 | 0.23 | 0.46 |
| Aluminum case<br>5x10x4 cm | 1 | 8.45 | 8.45 |
| switch | 2 | 0.87 | 1.74 |
| potentiometer 10K | 1 | 1.83 | 1.83 |
| **Total (US\$)** | | | **24.91** |
| Oscilloscope<br>Tektronix<br>TBS1072B | 1 |  | 800 |
| USB6218 | 1 | 1498 | 1498 |
| Labview | 1 | 2999 | 2999 |

The gain of the amplifier used was 99 and 1065 times calculated as:

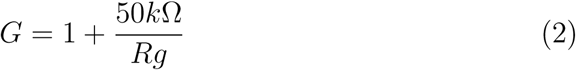

Where Rg is the external resistance to be modified with 511 ohms and 47 ohms to have the gains described.

The signal-to-noise ratio was calculated as:

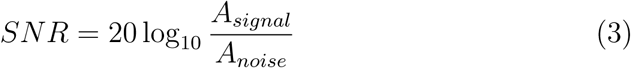

Where *A*_*signal*_ and *A*_*noise*_ are the RMS (root mean square) for each respectively.

The amplification system and the data acquisition hardware were placed inside a Faraday cage connected to the ground to avoid noise.

For de data acquisition and electrical stimulation, USB-6218 digital-analog converter with 250 KS/s (kilo-samples by seconds) and Labview software from National Instruments was used. The Labview program is based on graphical programming and consisting of a viewing window (front panel) and a programming window (block diagram). A A pulse generation program was made, and this consisted of a pulse with 100 *µ*s of duration with an amplitude ranging from 0-10 Volts at 1s intervals as described in Fig.3. A data acquisition program was made as described in a tutorial of the national instrument [13]. The rate acquisition was at 50kHz and stored in. lvm format (Labview measurement) and finally imported and analyzed using the Excel program.

**Fig.3.**
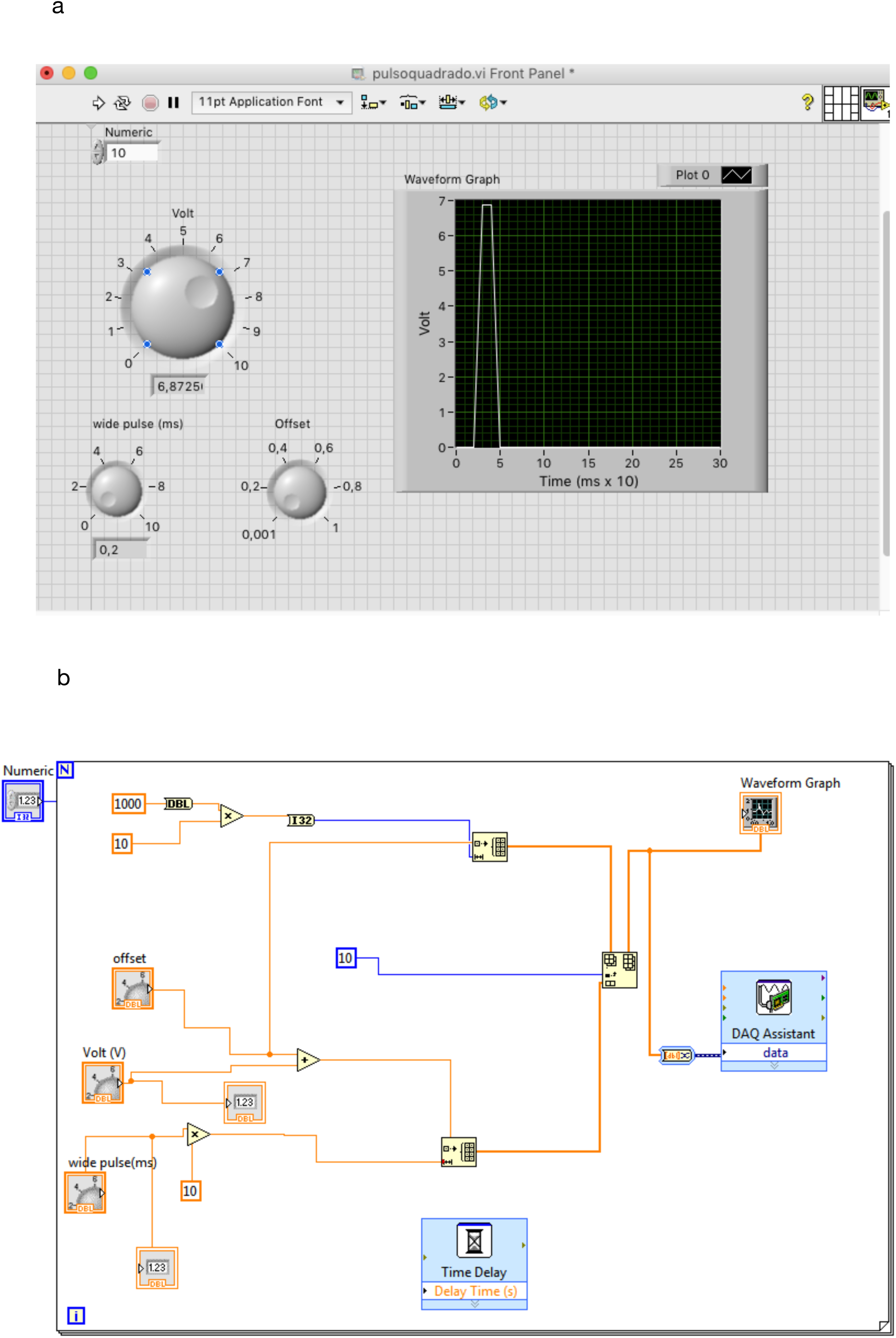
Front panel (a) and Block diagram (b) from Labview software for generating the stimulus pulse using the USB-6218.

## 3. Results and Discussion

### 3.1 Assembly of the experimental system

The amplification circuit was assembled as described in materials and methods. This showed stability in response and high signal to noise ratio of 25 dB avoiding the use of a filter for the purpose under study. The total cost of the amplification system was approximately US$ 25 (see Table 1), having a lower value compared to other commercial systems for the same purpose The most costly was the oscilloscope (US$ 800), the LabView program (US$ 2990) and the digital-analog converter USB-6218 (US$ 1498).

Over the past decade, several authors[6],[7],[8],[9],[14] have provided relatively low cost solutions using commercial equipment to amplify, filter and acquire data as well as the stimulation systems. Some of them use pin approach without having an invasive system in the AP registry but even anesthetized worms move slightly increasing the noise and varying the baseline making it very difficult to adjust the offset of the amplified signal. Low-cost systems are usually related to low acquisition resolution and noise issues. The proposed system considerably reduced the cost of amplification didn’t use filters but didn’t escape the use of high-cost systems such as oscilloscope and data acquisition system. The stimulation system has been simplified using the same acquisition system but this stimulus can also be generated using other platforms such as Arduino with limitation of the output voltage of the equipment to 5 volt.

### 3.2 Recording action potentials

After suctioning the nerve with the electrode suction and turning on the amplifier, it is possible to observe spontaneous fluctuations of potential (Fig. 4a), being these records were reproducible when the earthworm is slightly anesthetized (<6 min) and the suction electrode has suctioned the nerve cord properly. This means that the inner diameter of the pipette must be similar to the nerve diameter to increase the resistance between the internal and external Ag/AgCl electrodes to observe a potential variation.

**Fig. 4.**
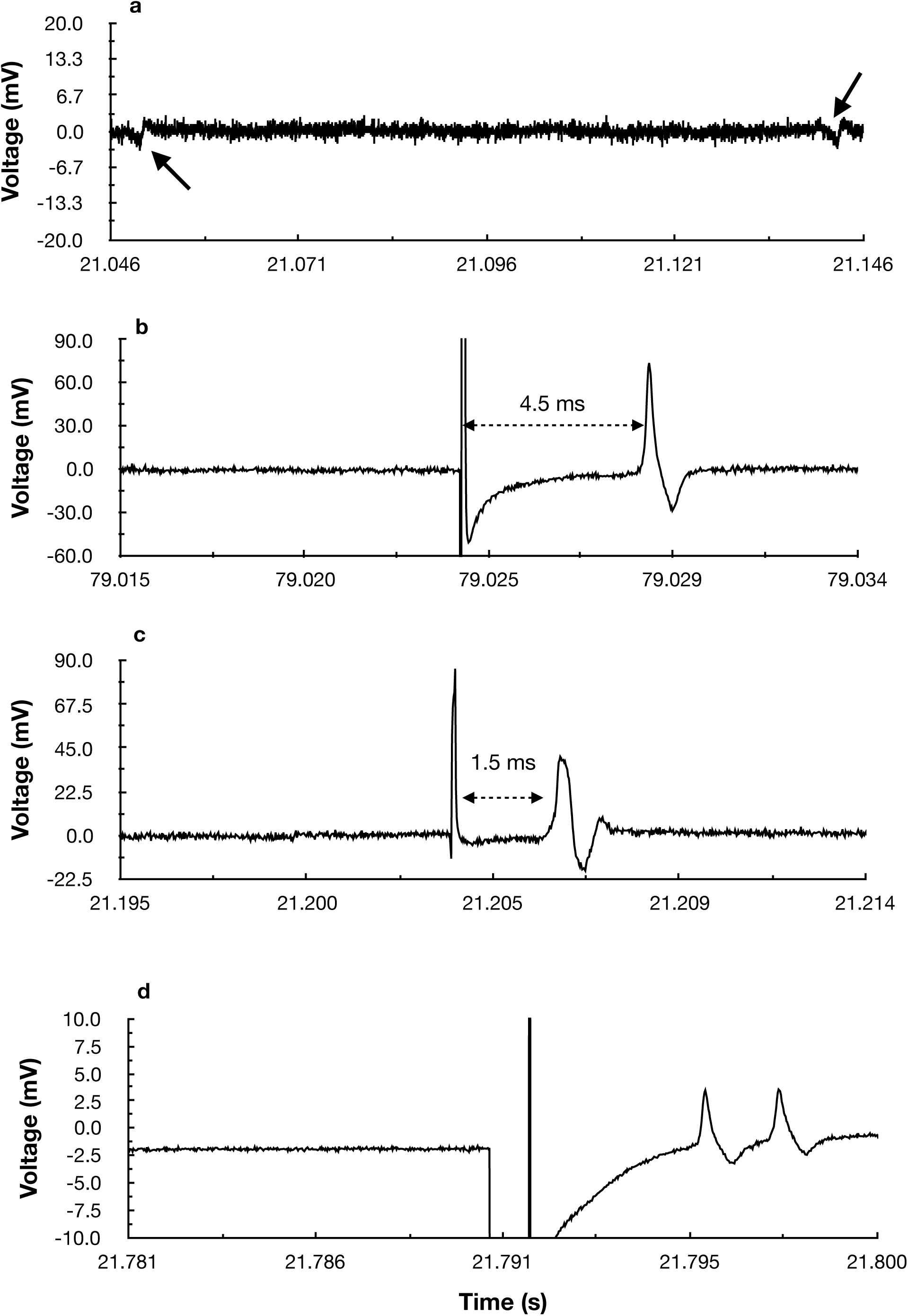
Action potential record from a ventral neural cord in the earthworm. a) spontaneous fluctuation of AP (indicated by continuous arrow) without electrical stimulus; b) characteristic AP lateral giant fiber with a latency of 5 ms; b) characteristic AP medium giant fiber with latency of 2 ms; d) Double AP by increased the stimulus duration from 100 us to 1 ms indicating the activation of both fibers in the nerve cord.

Under electrical stimulation conditions, there are characteristic fluctuations described as a development of action potential, with a pronounced depolarizing event followed a repolarizing face. It was possible to record action potential from lateral giant fiber (LGF) as medial giant fiber (MGF), featured for its conduction time with approximately 4.5 ms (Fig.4b) and 1.5 ms respectively (Fig.4c). This difference can be explained as described by Gunther at.al., 1961 and Shannon et al. (2014), increasing the diameter of the axon, increase their conduction velocity in a relationship that is the square root of the axon diameter. The diameter of the giant fiber is 0.05 mm for the LGF and 0.07 mm for the MGF approximately.

The peak-to-peak amplitude was approximately 100 mV (Gain = 100) (Fig.4b) and 60 mV (Fig.4c) but these values depend on the resistance between the reference electrode and recording electrode in the suction pipette because the measured value is the potential drop in external resistance. The conduction velocity was of 11 m/s for LGF and 33 m/s for MGF. The value found is similar to found by Rushton et al., (1944), Gunther et.al., (1961), kadt et al., (2010) and Shannon et al. (2014) and far from reported by Castelfranco and Hartline, 2016. According to Shannon et al. (2014), an increase of 1.4 times the conduction velocity between LGF and MGF is expected but it depends on the fiber size values which may vary depending on the earthworm analyzed.

Doubles AP was also observed when the ventral nerve cord was stimulated with a greater potential (from 1 to 9 V) or when the duration of the pulse increased from 100 *µ*s to 1 ms (Fig.4d). This is because the stimulation to voltage or pulse duration is high enough to stimulate both fibers in the nerve cord (MGF and LGF).

### 3.3 Troubleshooting

Some problems were encountered while registering AP in the neural cord. If the earthworm is anesthetized for a time larger than 6 minutes we don’t observe elicit spike, a situation similar to that described by Shannon et al., 2014. We recommend not to exceed 5 min.

AP logging is most often out of the baseline which can be offset by the offset regulator implemented in the amplifier.

If the pipette tip on the suction electrode has a diameter larger than the neural cord, AP may not be observed. The pipette tip must have a diameter similar to the neural cord. Failure to adhere properly greatly decreases the resistance between the electrodes and AP cannot be observed.

A system of articulated metal bars was used as a guide to the suction electrode, but a good approximation of the electrode to the neural cord is difficult. The use of a micro-manipulator system is necessary but it has a high cost.

### 3.4 Educational considerations

The work in the educational context resumes an experimental perspective that is losing itself every day with access to information internet way. The student wants quick answers through the use of sources and programs of simulation generating cognitive situations with prejudices without an adequate theoretical foundation. The described experimental system allowed the student to learn concepts of basic neurobiology in a ludic manner and stimulates him to the implementation of a new low-cost methodology for study pharmacological effects in AP and implementation of other techniques such as voltage-clamp current analysis, among others.

Therefore, practices such as these help in teaching-learning, leading to not only undergraduate courses but primary and secondary schools, stimulating students curiosity about basic processes in neurophysiology.

## 4. Conclusions

The present work allows recovering in a simple form the classic study of the propagation of nerve impulse by recording action potentials using components of low cost and simple to assemble and experience one of the basic foundations of neuroscience.

## Acknowledgements

We thank Professor Francisco Bezanilla Ph.D. for his valuable help and advice in assembling the circuit and experimental system.

## References

[1] A. L. Hodgkin, A. F. Huxley, Action potentials recorded from inside a nerve fibre, Nature 144 (3651) (1939) 710.

[2] W. A. H. Rushton, Action potentials from the isolated nerve cord of the earthworm, Proc. R. Soc. Lond. B 132 (869) (1945) 423–437.

[3] T. H. Bullock, Functional organization of the giant fiber system of lumbricus, Journal of Neurophysiology 8 (1) (1945) 55–71.

[4] P. Mill, Recent developments in earthworm neurobiology, Comparative Biochemistry and Physiology Part A: Physiology 73 (4) (1982) 641–661.

[5] J. Günther, Impulse conduction in the myelinated giant fibers of the earthworm. structure and function of the dorsal nodes in the median giant fiber, Journal of Comparative Neurology 168 (4) (1976) 505–531.

[6] R. Bähring, C. K. Bauer, Easy method to examine single nerve fiber excitability and conduction parameters using intact nonanesthetized earthworms, Advances in physiology education 38 (3) (2014) 253–264.

[7] K. M. Shannon, G. J. Gage, A. Jankovic, W. J. Wilson, T. C. Marzullo, Portable conduction velocity experiments using earthworms for the college and high school neuroscience teaching laboratory, Advances in physiology education 38 (1) (2014) 62–70.

[8] N. Kladt, U. Hanslik, H.-G. Heinzel, Teaching basic neurophysiology using intact earthworms, Journal of Undergraduate Neuroscience Education 9 (1) (2010) A20.

[9] O. R.F., Using the daq assistant to automatically generate labview code (2019). URL http://www.science.smith.edu/departments/neurosci/courses/bio330/labs/L4giants.html

[10] M. Roberts, The giant fibre reflex of the earthworm, lumbricus terrestris, L, J. Exp. Biol 39 (1962) 229–237.

[11] A. P. M. Lockwood, “ringer”, solutions and some notes on the physiological basis of their ionic composition, Comparative biochemistry and physiology 2 (4) (1961) 241–289.

[12] B. R. Johnson, S. A. Hauptman, R. H. Bonow, Construction of a simple suction electrode for extracellular recording and stimulation, Journal of undergraduate neuroscience education 6 (1) (2007) A21.

[13] N. Instrument, Using the daq assistant to automatically generate labview code (2019). URL http://www.ni.com/tutorial/4656/en/

[14] A. Gonzalez-Perez, R. Budvytyte, L. D. Mosgaard, S. Nissen, T. Heimburg, Penetration of action potentials during collision in the median and lateral giant axons of invertebrates, Physical Review X 4 (3) (2014) 031047.

[15] A. M. Castelfranco, D. K. Hartline, Evolution of rapid nerve conduction, Brain research 1641 (2016) 11–33.

